# QCM: real-time quantitative quality control of single-molecule localization microscopy acquisitions

**DOI:** 10.1101/2024.07.23.604731

**Authors:** Sébastien Mailfert, Meriem Djendli, Roxane Fabre, Didier Marguet, Nicolas Bertaux

## Abstract

Single molecule localization microscopy (SMLM) has revolutionized the understanding of cellular organization by reconstructing informative images with quantifiable spatial distributions of molecules far beyond the optical diffraction limit. Much effort has been devoted to optimizing localization accuracy. Among them, assessing the quality of SMLM data in real-time, rather than after lengthy post-acquisition analysis, represent a computational challenge.

Here, we overcome this difficulty by implementing an innovative mathematical approach to drastically reduce the computational analysis of particle localization. We have therefore designed the Quality Control Map (QCM) workflow to process data at a much higher rate than that limited by the frequency required by current cameras. Moreover, QCM requires no parameters other than the PSF radius characteristic of the optical system and only a GPU card to reach its computational speed. Thus, QCM is robust and adaptable to any type of input data. Finally, the QCM off-line mode can be used to evaluate synthetic or previously acquired data, and as a tool for teaching the basic concepts of the SMLM approach.

**Teaser:** QCM, a parameter-free algorithm, calculates indicators for instant feedback on single-molecule localization precision experiments

## INTRODUCTION

In system biology, the combination of “omic” approaches can benefit significantly from Smart Microscopy (SM) to bridge the gap between cellular events and organism-level phenomena, enabling the unravelling of complex biological networks (*1*). By providing key spatio-temporal observables, photonic microscopy has become the cornerstone of scientific research in biology, to which SM is giving it a new technological breath (*2*). Innovative SM approach combines cutting-edge hardware, sophisticated software and powerful algorithms to facilitate the use of increasingly complex microscope modalities. As anticipated, SM offers the possibility of combine imaging procedures thanks to automated data acquisition in a single experiment, in a simplified and reproducible way. As the amount of information increases, approaches based on real-time analysis or machine learning algorithms enable acquisition parameters to be adjusted on the fly (*3*) and/or large quantities of data to be processed to identify patterns, anomalies and subtle changes, ultimately enabling autonomous decision-making or rapid and accurate diagnosis (*4*). Thus, next-generation microscopes are poised to assist humans in automating the acquisition and analysis of data in regions of interest driven by specific events. To this purpose, it is necessary to provide real-time feedback to adjust parameters, optimize imaging conditions and dynamically explore samples.

This requirement is particularly relevant to photonic microscopy approaches based on single-molecule localization microscopy (SMLM), which has revolutionized the understanding of cellular organization by reconstructing informative images at the nanoscale (*5-8*). SMLM observations can inherently produce well-resolved images from which biologically relevant information can be determined such as the nanostructure and stoichiometry of macromolecular complexes (*9*), provided that the SMLM data production process is properly mastered to resolve a given biological question (*10, 11*). In this respect, many efforts have been made to optimize not only the sample preparation (unbiased fixation, labeling procedures, etc.) (*12-16*), but also the acquisition modalities (laser power, camera integration time, stabilized optical systems etc.) (*17-20*) or the design of dedicated quantitative analytical methods (*21-23*).

The overall process of generating SMLM data, which includes image acquisition, handling and analysis, is time-consuming, and results are highly dependent on the quality of the data acquired to achieve a given localization accuracy and to avoid misleading interpretations. Therefore, a computational challenge is to estimate this localization accuracy before rather than after data acquisition to save time and avoid losing valuable samples (Fig. 1a). Still, most of software packages ensure robust quantitative *a posteriori* analysis (see for review (*24, 25*)), assuming that the data have been recorded appropriately for reconstructing super-resolution images, given that no further adjustment or correction of the acquisition parameters can be made like NanoJ-SQUIRREL (*26*) or SuperStructure (*27*). For instance, the former provides a quantitative assessment of SMLM results by generating a quantitative evaluation of super-resolution images to help experimenters optimize imaging parameters; this approach is based on comparing diffraction-limited images and super-resolution equivalents.

**Fig. 1.**
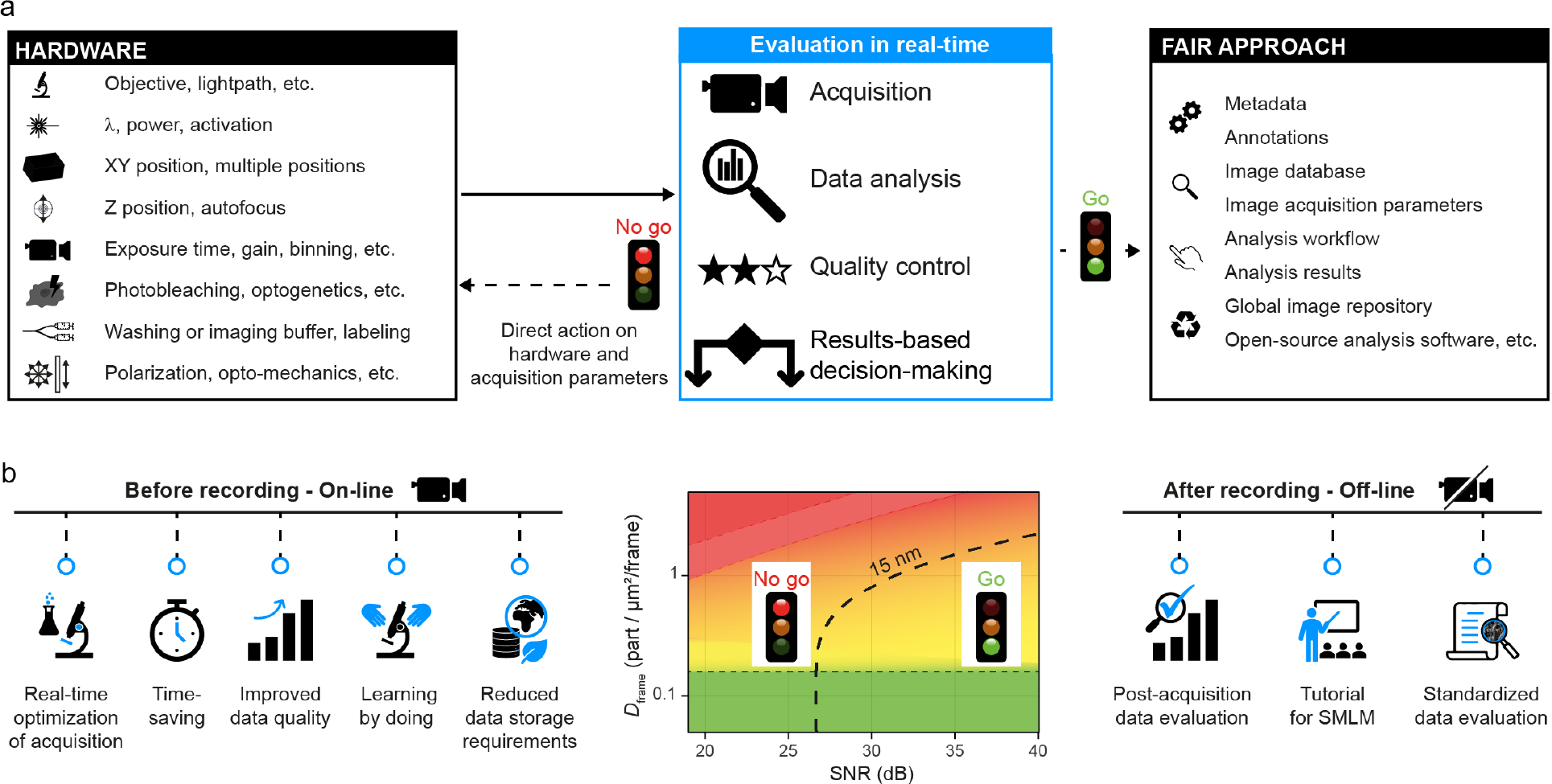
A need for comprehensive quality control tools for SMLM acquisitions. (**a**) Smart microscopy guidelines aim to integrate quality control tools from the earliest steps of the SMLM acquisition process up to post-process analysis. (**b**) The density-SNR space diagram (middle panel) summarizes the expected localization accuracy as a function of the two key indicators, SNR and *D*_frame_. The black dashed line marks the limit for achieving an overall particle localization accuracy of e.g. 15 nm (*35*). The on-line mode of QCM processes these key indicators in real-time, providing the instant feedback needed to optimize acquisition parameters prior to data recording (left panel). In addition, the use of the off-line mode in post-acquisition data analysis provides a tool for standardized data review or for teaching SMLM methods (right panel).

Consequently, analytical tools for assessing the quality and robustness of SMLM data at any time such as those enabling *a priori* quantitative control of data, are therefore in high demand from a broad community of cell biologists (Fig. 1a). The aim is to carry out analyses in real time in order to adjust the acquisition parameters for optimal data recording. This approach should make it possible to avoid time-consuming and unnecessary data acquisition, when a posteriori analysis will reveal only poor-quality and misleading data. Some strategies have implemented new computational strategies to speed up image acquisition or processing (*28, 29*). Computer architecture design is another means of achieving high computing performance (*26, 28, 30-33*). For instance, QC-STORM (*30*), a GPU-based software package, performs real-time image processing and generates a list of particle localization, but lacks precise quantification, relies only on indicators on the full dataset and provides only histograms. Another method computes the Fourier Ring Correlation measurement in real-time (*34*).

Alternatively, hardware developments have been implemented to compute multi-emitter fitting in real-time (*32*).

Considering that the expected localization accuracy is directly dependent on two parameters - the signal-to-noise ratio (SNR) and the particle density per frame (*D*_frame_) - we have implemented the Quality-Control Maps (QCM), a parameter-free algorithm that represents a major advance in the SMLM field and extends the SPT and SMLM algorithms previously developed (*35, 36*). Here, by implement an original mathematical approach, we were been able to harness the computing power of conventional computers to carry out the analysis of 2048 × 2048 pixels images at a rate of over 100 frames/second, which is sufficient for real-time analysis.

The special feature of QCM is that it displays in real-time quantitative maps and histograms of local (zoomed-in areas) and global (full frame) of a set of indicators to assess the quality of SMLM data in an easily understandable way thanks to its color coding. These include the PSF size in xy, and xyz positions, signal-to-noise ratio (SNR), background, intensity, and precision of localization. It should be noted that another major advance of QCM relies on the SNR (in dB) as the most relevant contrast parameter for summarizing expected achievable precision; indeed, the root mean square of the localization precision when expressed by SNR depends weakly on noise model or density/frame (see in Mailfert et al. (*35*) the Materials and Methods section and Fig. S1).

Thus, the workflow of the Quality-Control Maps (QCM) software has been designed to conduct real-time analysis of data, providing users with key observables in the decision-making process. If the results do not meet predetermined criteria such as a given accuracy of molecular localization, users can intervene on the setup, ensuring optimal acquisitions in line with the FAIR principles (*37*). Used prior to acquisition (Fig. 1b, left panel), QCM primarily saves time, improves data relevance and reduces data storage requirements. In addition, the use of QCM for post-acquisition data analysis provides a standardized tool for educational purposes or for peer review of data (Fig. 1b, right panel).

## RESULTS

### The QCM heuristic

The QCM algorithm is divided into two main modules, Ultra-Fast Unsupervised Localization (UFUL) and Quality Control (QC) (Fig. 2). The first one is based on an innovative mathematical approach designed to drastically reduce the computational steps involved in particle localization analysis. As a result, UFUL performs the particle detection/localization steps at a much faster rate than the frame rate acquisition by standard cameras used in SMLM. The second module then uses the UFUL results to estimate the relevant SMLM indicators in real-time, and displays their histogram distributions and map representations.

**Fig. 2.**
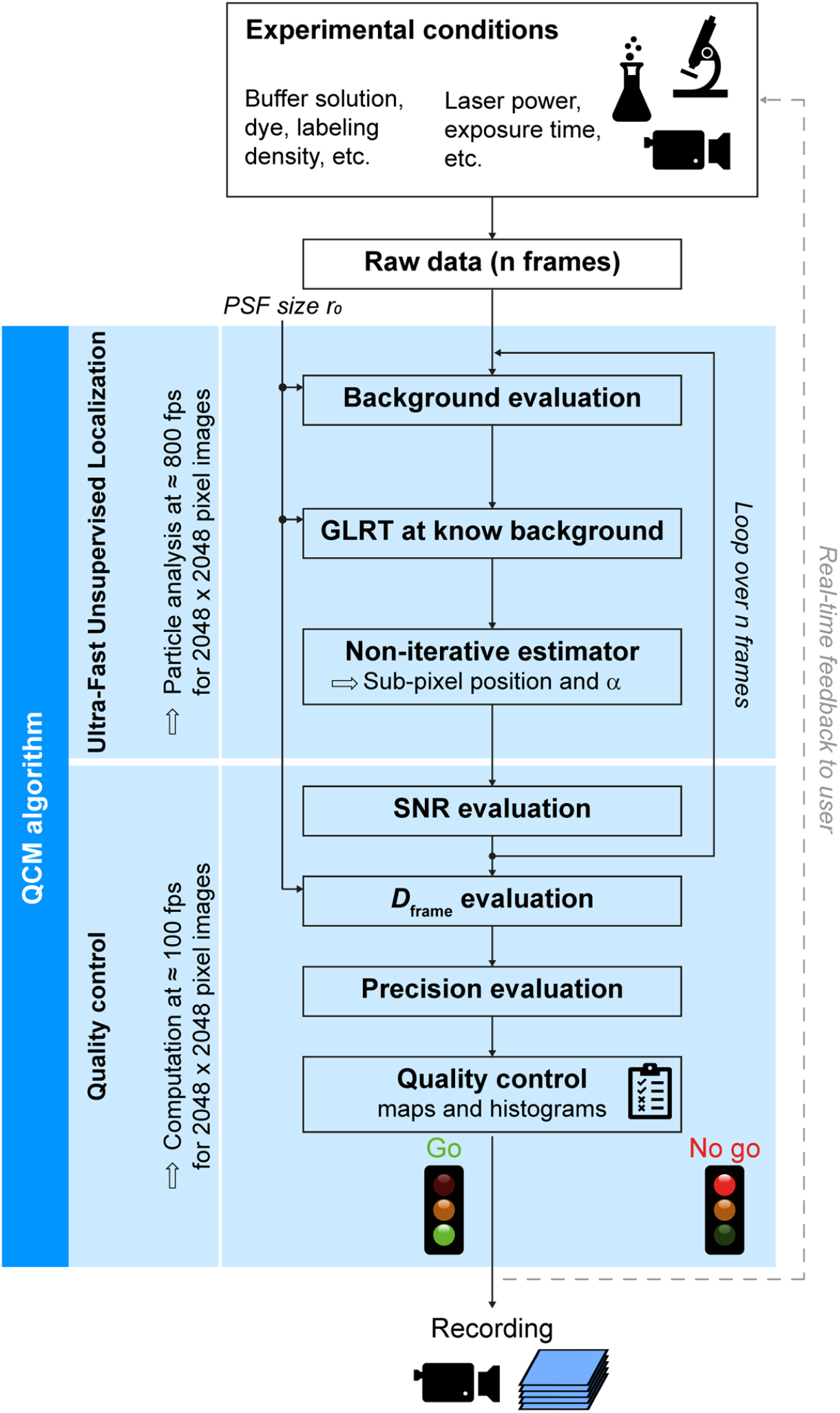
QCM workflow. The QCM algorithm extends from the initial setting of microscope parameters to the decision whether or not to record SMLM data. It combines (1) the Ultra-Fast Unsupervised Localization (UFUL) algorithm to perform the particle detection/localization steps at a rate of ≈ 800 fps for 2048 × 2048-pixel images, i.e. at a speed higher than that of image acquisition by current SMLM cameras, with (2) the Quality Control (QC) module for real-time estimation of indicators: *D*_frame_, SNR, and localization accuracy. Acquisition parameters that pass quality control criteria are used to start recording data.

Classically, single particles localization involves a detection step and then an estimation at high resolution (*i*.*e*. sub-pixel) of the particle position in the (*i, j*) plane and the size of the point spread function (PSF) on the *i* and *j* axes (*36*). To solve this problem, the regular procedure is based on a maximum likelihood estimator (MLE) or a minimum mean square error (MMSE) estimator. The main objective is to avoid using estimators on a region of interest (ROI) devoid of particles.

Within this framework, we have previously provided mathematical developments optimizing the detection step by implementing a generalized likelihood ratio test (GLRT) at known background (*35*), which means that the mean *m* and variance *σ*^2^ of the background are known. This test is an effective unsupervised detection tool whose threshold is set by the probability of false alarm (*PFA*) (*38-40*); it is primarily designed as a detector in a ROI with a working window of dimension *ω*, for the absence (*H*_0_ hypothesis) or presence (*H*_1_ hypothesis) of a particle. The GLRT can also be conceived as an estimator since, under the *H*_1_ hypothesis assumption, it builds the intensity image using an estimator in the sense of the MMSE estimator, which is close to the Cramer-Rao Bounds (CRB). As such, the GLRT performs an adaptive filter to carry out the estimation of the intensity as previously described (see for instance the Supplementary Note 2 in (*35*)). However, its application as initially conceived cannot handle the real-time data flow of SMLM acquisitions.

To overcome this difficulty, we have rewritten the mathematical operations of the GLRT detector at known background to considerably accelerate the computational steps, without impacting its robustness (see Materials and Methods, for details). Practically, this implies to evaluate first the mean 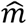 and variance 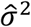 of the background as in (*35*). Then, the GLRT assess the presence of a signal at each pixel and when a signal is detected, the estimator searches the sub-pixel positions of the PSF. UFUL computes 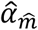 as the expression of the intensity 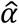 minus the image background 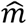. Three separable convolutions are processed for each pixel to estimate the background, its variance, and the signal intensity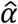, respectively. It is then possible to estimate the sub-pixel position of each detected particle and the size of its PSF. To do this, the PSF is modeled by a Gaussian function from which UFUL uses a logarithm of the intensity 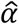 to derive literal expressions for estimating for each particle, its PSF radii *r*_*i*_, *r*_*j*_ and sub-pixel coordinates *i*_0_, *j*_0_ on axes *i* and *j*, respectively.

We test the UFUL performances to ascertain that the analyses coincide with the MMSE estimation, with respect to the variances in positions. This was done on realistic synthetic data, *i*.*e*. on data close to the levels of noise, signal, PSF size, etc. that are typically the ones observed on experimental SMLM. We report that UFUL overlap those of an MMSE estimator; both being close on the CRB. For a PFA of 10^−6^, the detection probability *PD* ≈ 100% for any *SNR* > 20 *dB* (Supplementary Text, Fig. S1 and S2).

Moreover, UFUL provides the estimation of *r*_0_ on both *i* and *j* axes with the estimator 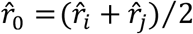 regardless of the PSF size and the working window. Consequently, when image acquisitions are performed with an astigmatic lens (*41*), the axial PSF of a particle is distorted in *i* and *j* axes of the focal plane as a function of the particle’s position on the optical axis, enabling it to be localized in 3D (Supplementary Text, Fig. S3). Under these conditions, the size of the working window *ω* for the GLRT detector set at 8 × 8, 10 × 10, 12 × 12, 14 × 14, or 16 × 16 pixels, depends on *r*_0_, the size of the PSF.

### UFUL computation rate performance

We evaluated the UFUL performances on synthetic datasets generated at a given density of particle per frame and for different image sizes (Fig. 3). The analyses were obtained on a computer with the option of processing data with CPU (central processing unit) or GPU (graphical processing units) processors (see Material & Methods for the specifications).

**Fig. 3.**
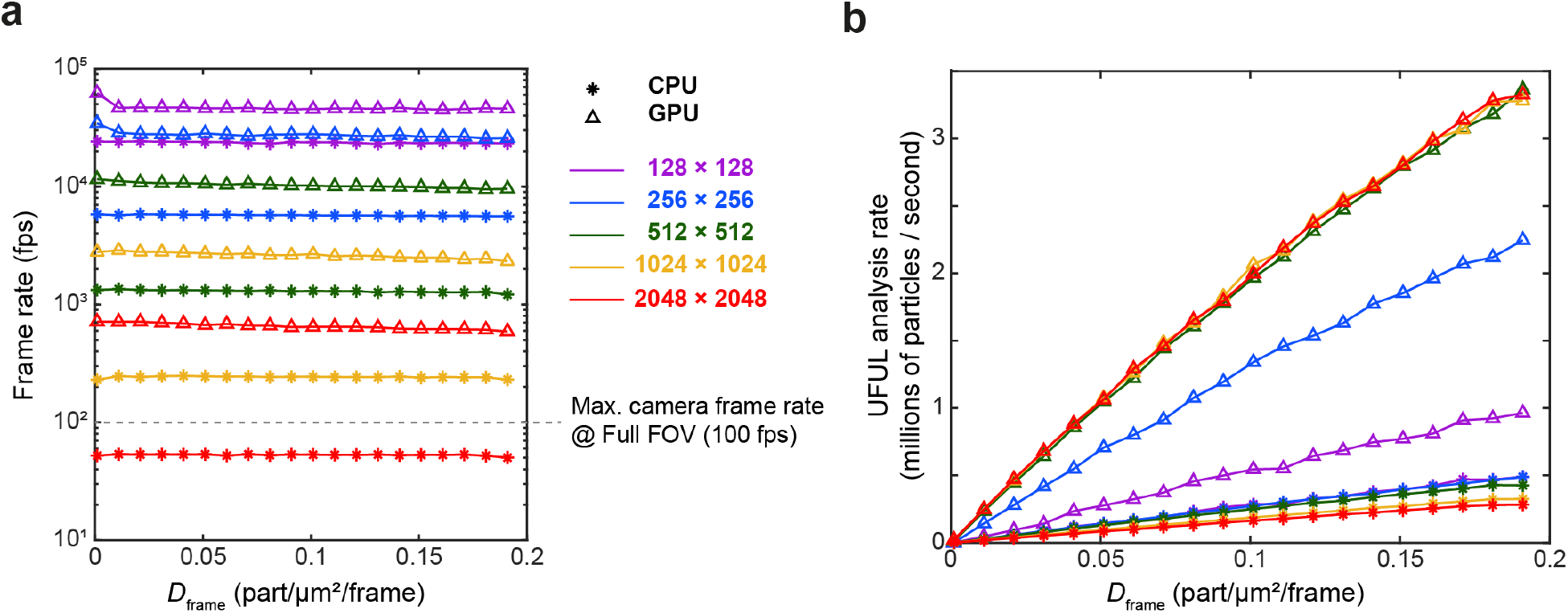
Comparison of CPU and GPU UFUL computation rates as a function of *D*_frame_ and image sizes on realistic simulated data. (**a**) Number of frames analyzed per second (fps) based on given *D*_frame_ values for images ranging from 128 × 128 to 2048 × 2048 pixels. (**b**) UFUL analysis rate expressed as number of particles detected and estimated position per second as a function of *D*_frame_ for different image sizes.

The performances are expressed in number of frames analyzed per second (Fig. 3a). The computation times correspond to the analysis of 16-bit raw images stored in the PC RAM, from which the detection/estimation process returns a list of particles with position, size of 2D or 3D PSF astigmatism, intensity, SNR, noise level, and position errors in the PC RAM. This time mainly results from the one used by the GLRT detector and therefore, for *D*_frame_ values ranging from 0 to 0.2 part/µm^2^/frame, the number of particles has hardly any impact on the performance of detection/estimations steps (Fig. 3a).

The UFUL module incorporates a computing segment specifically designed to optimize computation time to achieve a data flow of over 5 GB/s, enabling it to perform data analysis at a rate well above the acquisition performance of a standard SMLM camera, i.e., ≈ 100 fps for 2048 × 2048-pixel images. Indeed, with GPU, the analysis rate for particle detection and position estimation reaches over 10,000 fps for 256 × 256-pixel images and up to ≈ 800 fps for 2048 × 2048-pixel images (Fig. 3a). Furthermore, GPU computing increases the number of particles analyzed per second by a factor up to 10 compared to CPU computing on large images (Fig. 3b). For the GPU, images of size equal to or greater than 512 × 512 pixels present a similar number of particles analyzed per second, unlike the CPU where the cache miss is significant.

Overall, the GPU speeds up analysis considerably, with a more noticeable difference on large images compared to the CPU. It should be noted, however, that while the UFUL computation rate with a GPU is well above the frame rate of SMLM camera, performance with a CPU remains above this threshold for image up to 1024 × 1024-pixels (Fig. 3a). As a result, the second module of QCM can process the output of UFUL results and display the relevant quality control indicators in real time.

### QC module and the QCM interface

The QC module relies on UFUL results for the image background, particle positions, intensity and PSF size. It evaluates in real-time the key quality control indicators -*D*_frame_, SNR and precision of localization parameters – and displays using histograms and maps. Thus, we set up a graphical interface for easy, real-time evaluation of QCM analysis results (Fig. 4 and Supplementary Video 1). In the opening panel, QCM requires no parameterization other than the following physical parameters (Fig. 4a):

**Fig. 4.**
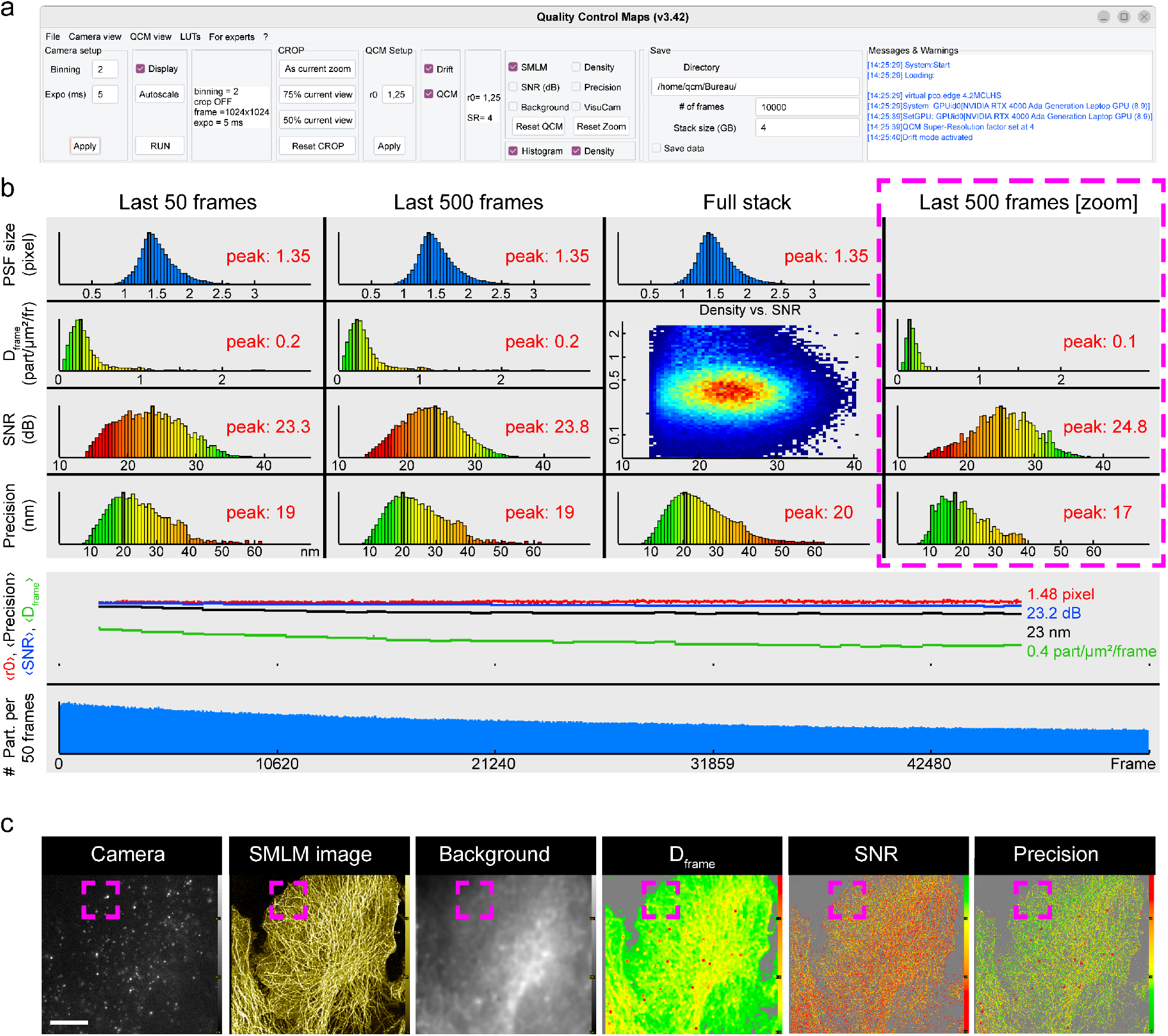
General description of the QCM graphical interface. dSTORM imaging of β-tubulin in COS-7. (**a**) Main functions from left to right: *camera setup*: binning and exposure time; *QCM setup*: setting the PSF size of the microscope; *Visualization & indicators*: histogram or map visualization options (see below (**b**) and (**c**)), selection of the QCM calculated parameters; *Save*: file and data acquisition saving options, and *messages & warnings*. (**b**) Real-time histograms of the PSF size *r*_0_ (pixels), *D*_frame_ (particles/µm^2^/frame), SNR (dB), precision (nm), and the number of particles detected per image can be displayed for the last 50, 500, or cumulative full field of view (FoV) frames or the last 500 frames on a zoomed ROI. Indicator values are also traced over time to assess their stability. (**c**) Real-time QCM windows – *Camera*: shows in real-time the full FoV or zoomed area of a frame recorded by the camera; *SMLM image*: compilation of detected particle localizations; *Background*: background intensity; *D*_frame_: color-coded *D*_frame_ values; *SNR*: color-coded SNR values; *Precision*: color-coded of the root mean square particle precision estimated from the combination of *D*_frame_ and SNR indicators. Scale bar: 20 µm and insert 5 µm on a side. For more details on the QCM display, see the user manual.

- The characteristic PSF radius (*r*_0_) of the optical system, a physical characteristic inherent in the optical system for a given excitation wavelength and numerical aperture of the objective. This value must be expressed in pixels. As part of internal quality control, QCM displays the *r*_0_ histogram evaluated during data acquisition.
- The binning and exposure time parameters of the camera.

QCM displays in real-time the histograms of the PSF radius (*r*_0_), *D*_frame_, SNR and precision parameters. Analyses are visualized on the last 50, 500 or full stack of images; it is also relevant for specific applications to estimate these indicators on a zoomed region of interest (Fig. 4b and Supplementary Video 2). Moreover, when a biological question requires achieving a given SMLM precision (e.g., dimensionality or count of macromolecular complexes, inter-distances between macromolecular structures, etc.), QCM offers the option of displaying the density-SNR space diagram in real-time allowing standardized evaluation of experimental data. Other options allow to display the images captured by the camera, the SMLM image reconstructed at the time being, or those of quality control indicators (background, *D*_frame_, SNR, and precision parameters) (Fig. 4c).

### Assessment of the robustness of QCM analyses

To assess the robustness of quality control indicators calculated by QCM, we collect stacks of 2,000 images of DNA origami nanorulers as nanoscopic benchmark structures (*42*). The SMLM DNA-PAINT imaging technique is used to assess the metrological traceability of nanorulers with marks 80 nm apart. The data acquired at different laser powers and camera integration times were analyzed in real-time by QCM. Each initial acquisition is short, around one minute per condition, but long enough to display informative SNR and *D*_frame_ histograms for deciding whether or not to continue data acquisition. QCM results were compared with those obtained on the same dataset using UNLOC and GATTAnalysis as post-acquisition analysis tools (Fig. 5). As illustrated, the images reconstructed by UNLOC provides a qualitative estimate of the nanorulers while GATTAnalysis evaluation is based on three parameters, the pass ratio, i.e. the percentage of good spots, the mark-to-mark distance in nm and the fraction of nanorulers at a precision threshold better than 20 nm. For example, under acquisition at a laser power of 37 mW and a camera integration time of 36 ms, the histograms of *D*_frame_ and SNR provided by QCM peak at 1.3 particles/µm^2^/frame and 22.4 dB, respectively. Under such conditions, we cannot expect to ensure robust SMLM resolution, as assessed by post-acquisition analyses, where only 12% of nanorulers achieve accuracy better than 20 nm. A go/no-go decision based on the SNR and *D*_frame_ histogram distribution provided by QCM analyses on a small number of frames is in good agreement with the results of post-acquisition analyses provided by the UNLOC or GATTAnalysis algorithms. Therefore, QCM enables instant adjustment of camera integration time and/or laser power to find the optimal acquisition parameters for achieving the highest possible localization accuracy on the samples, before starting the SMLM data acquisition.

**Fig. 5.**
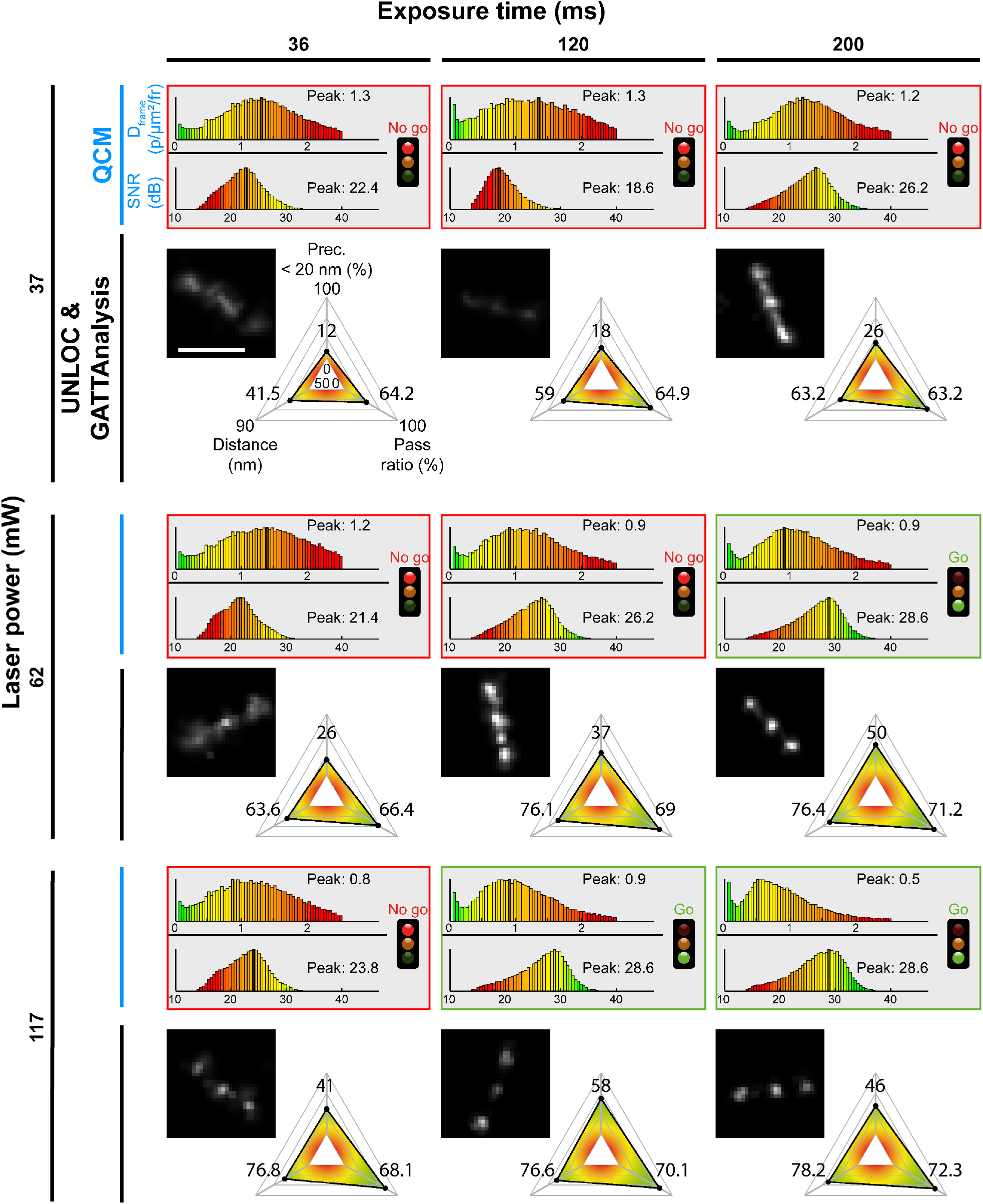
Validation of the robustness of QCM analyses. Image stacks of 2,000 frames of DNA origami 80 nm nanorulers were acquired at different camera integration times and laser powers. Data analyzed by QCM (SNR and *D*_frame_ histograms) were compared with the color-coded spider plots quantified by GATTAnalysis software and the UNLOC results (scale bars: 160 nm) with data at a precision threshold better than 20 nm. The real-time go/no-go decisions based on QCM analyses are in good agreement with the post-acquisition analyses.

### QCM-optimized SMLM acquisition on biological samples

Next, we test QCM for imaging biological samples using different SMLM methods, to investigate the robustness of the displayed key parameters, SNR and *D*_frame_ histograms, before starting data recording. To do this, we compared the QCM results obtained on the first stack of 2,000 images with those obtained by post-analysis with UNLOC on the whole recorded dataset (up to 50,000 frames).

Cellular expression of β-tubulin and the nuclear pore protein Nup133 was imaged by dSTORM SMLM (Fig. 6). After chemical fixation, cells were incubated with primary antibodies before staining detection with fluorescently-labelled secondary antibodies. Samples were imaged in freshly prepared dSTORM buffer, and laser power and camera integration time were adjusted to image β-tubulin in COS-7 cells and Nup133 in HeLa cells respectively. Since QCM quality control on given imaging conditions prior to acquisition can be based on a few hundred frames, it is fast enough to avoid distorting the recording of a whole dataset due to lengthy adjustment procedures (e.g., due to photobleaching or dSTORM buffer deterioration) (Supplementary Video 3). For example, the initial QCM analyses of β-tubulin imaging in COS-7 cells were carried out on just 2,000 frames displaying informative and robust SNR and *D*_frame_ histograms (Fig. 6a). Besides, the QCM analyses can be operated for the entire duration of the data recording, so that the mean values of indicators are tracked over time, enabling their stability or inconsistency to be assessed, for instance in the event of focal plane loss (Fig.4b). Finally, QCM and UNLOC analyses carried out on the same number of frames show that 35% and 49% of detected signals have an estimated precision greater than or equal to 20 nm for β-tubulin and Nup 133, respectively (Fig. 6b).

**Fig. 6.**
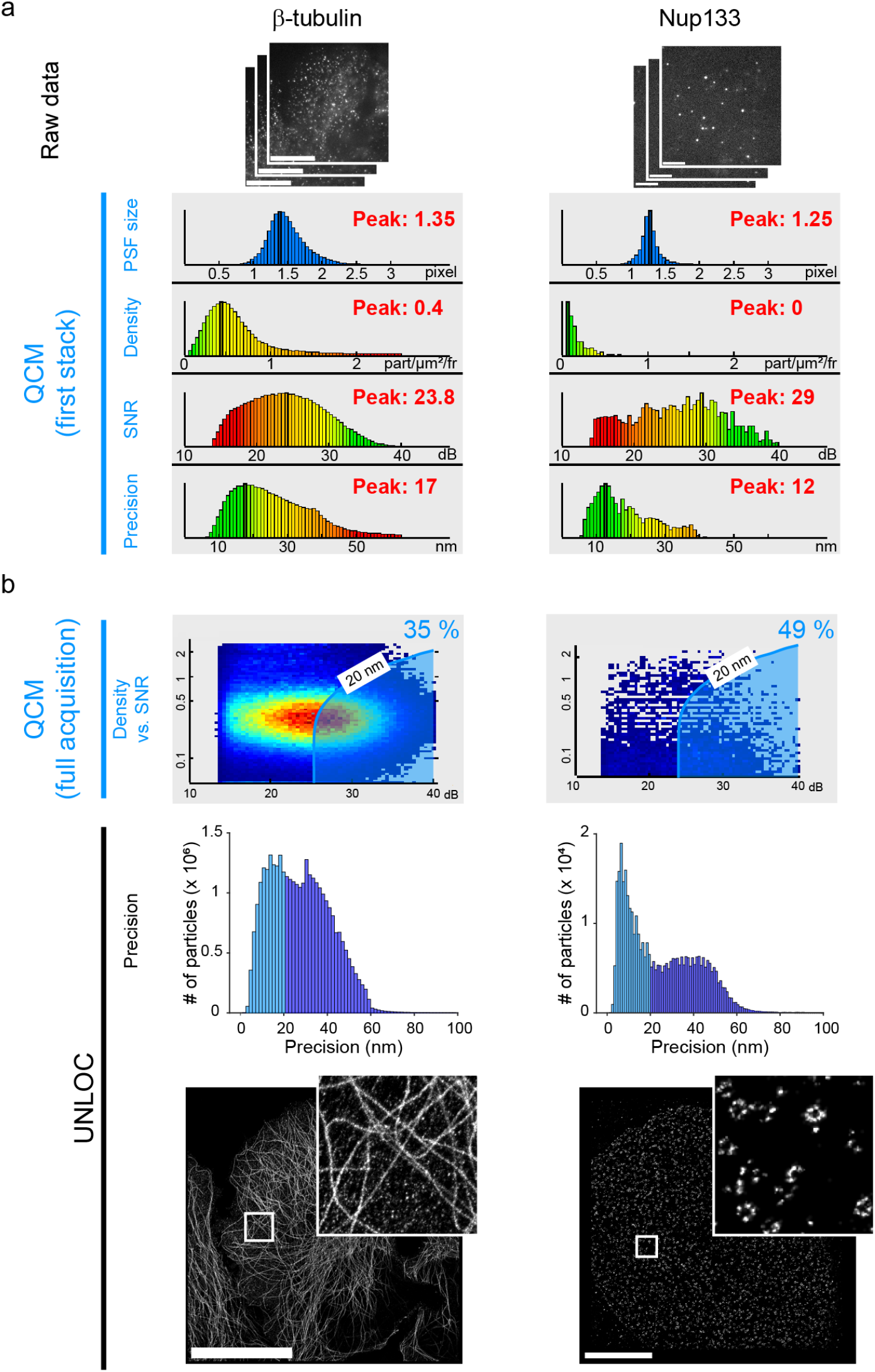
QCM-optimized dSTORM acquisitions. dSTORM imaging of β-tubulin and Nup133 labelling in COS-7 and HeLa cells, respectively. (a) QCM histograms and maps were computed from the first 2,000 frames. (b) Post-acquisition analysis of the recorded raw data sets. The density-SNR space diagrams displayed by QCM for β-tubulin and Nup133, reveal that 36% and 49% of detected particles have a localization precision better than 20 nm, respectively. UNLOC show the integrated Gaussian reconstructed images for particles with precision better than 20 nm. Scale bars: 20 µm (inserts: 5 µm on a side) and 5 µm (insert: 1.5 µm on a side) for β-tubulin and Nup133, respectively.

As discussed in the Supplementary Text and Fig. S4, the detection/estimation achieved by UFUL is primarily designed for image analysis under low density condtions, i.e. *D*_frame_ less than ≈ 0.2 part./µm^2^/frame. To overcome this limitation, a density evaluation calibration has been integrated so that the algorithm returns realistic density values. But the fine analysis of raw data in most cases requires post-acquistion analysis with a dedicated algorithm based on heuristic for reliable particle localization at variable local density. Nevertheless, the analysis obtained with QCM give a very good estimate of those obtained with a dedicated algorithm such as UNLOC (Fig. 6b).

Among other SMLM methods, DNA-PAINT, based on the transient association of a fluorescently-labeled probe with a target molecule, has become particularly popular due to the ability to adjust experimental conditions to the expression level of the proteins of interest being visualized. The signal detection is mediated by pairing a docking oligonucleotide coupled to a target probe that recognize a protein of interest with an imager, i.e., a fluorescently labelled complementary oligonucleotide freely diffusing in the buffer (*43*). This method relies on the concentration of imager to control the density of transient docker/imager hybridization per frame enabling a stochastic detection of the protein of interest by recoding fluorescence signals.

For two-color DNA-PAINT experiments (Fig. 7), we used an automated workflow system to deliver sequentially into a channel slide, the respective imagers to detect in HeLa cells the mitochondrial 20 kDa outer membrane protein TOM20 and the major building block of microtubules α-tubulin (see Materials and Methods). For acquisitions at appropriate D_frame_, the imager concentrations were pre-adjusted using QCM over just 500 frames, i.e. an acquisition time of 18 s, as shown in Fig. 7b to define the conditions required for TOM20 protein imaging. QCM analyses were performed at three successive imager concentrations on the same sample preparation. At 1.5 nM, the precision of localization was significantly impaired. At 0.3 nM, the QCM returns significant intracellular variability in D_frame_ and SNR values, with poorly resolved area as illustrated in the insert. At a 10-fold lower imager concentration, i.e. 30 pM, QCM returns adequate precision of TOM20 localization for any intracellular area. The relevance of the QCM analyses obtained on a few hundred images is demonstrated by the comparison with the reconstructed images obtained with UNLOC on complete datasets. We thus detected the distribution of TOM20 and α-tubulin in HeLa cells. Consequently, the efficiency of QCM makes it possible to adjust experimental conditions in real time for optimal DNA-PAINT-based multicolor sequential localization of multiple cellular components such as TOM20 and α-tubulin, as illustrated by the integrated-Gaussian reconstructed images from the post-processed UNLOC analyses (Fig. 7b). Thus, QCM is a key asset for unlocking the power of the multicolor DNA-PAINT SMLM approaches.

**Fig. 7.**
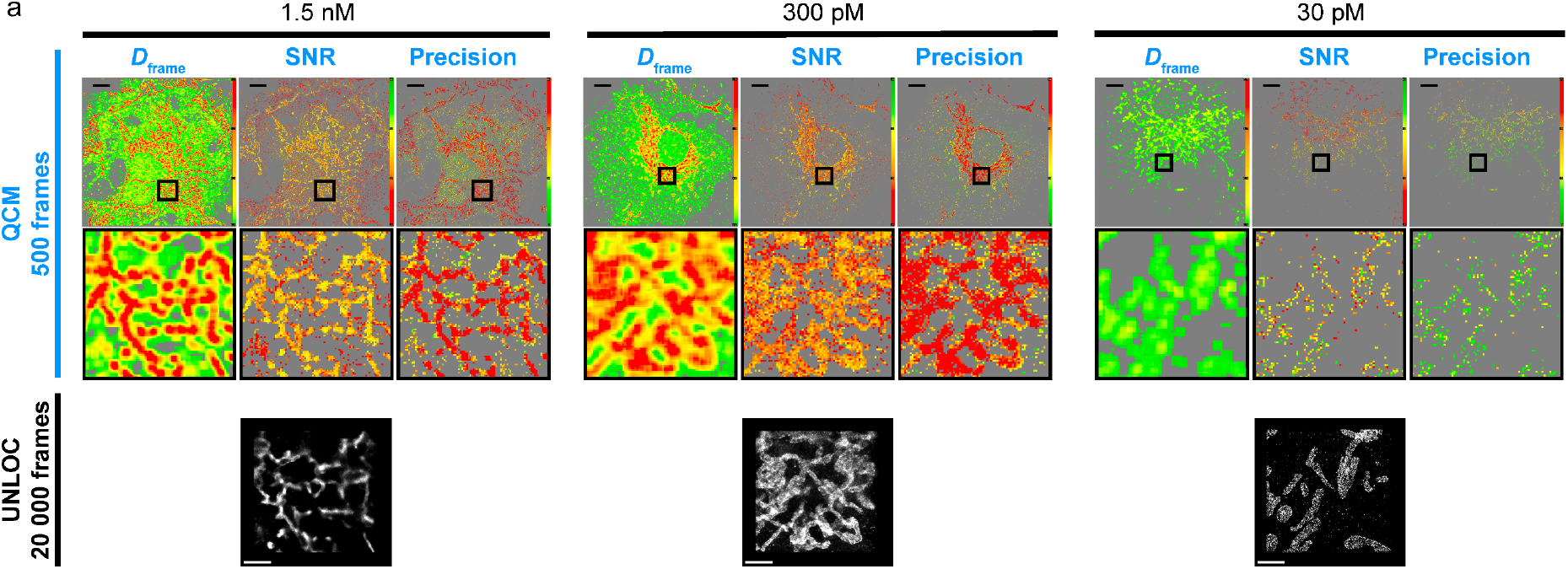
QCM -optimized acquisitions of two-color DNA-PAINT data. (**a**) Detection of TOM20 in HeLa cells by DNA-PAINT imaging on 500 frames. Real-time QCM analyses at different imager concentrations anticipates incorrect and inappropriate acquisitions based on poor *D*_frame_, SNR, and precision indicator values at imager concentrations above 30 pM. Post-acquisition analyses validate the quality control observations, as evidenced by the shape of the mitochondrial network in the reconstructed images by UNLOC. Scale bars: 10 µm (in the inserts: 1 µm). (**b**) Reconstructed images from post-processed α-tubulin and TOM20 data with UNLOC. The concentrations of imagers for two-color DNA-PAINT acquisition with sequential fluid exchange were pre-adjusted with QCM. Scale bars: 20 µm (in the insert: 2 µm).

## DISCUSSION

In line with FAIR principles (*37*), the emergence of smart microscopes where tools such as QCM enable quantitative data analysis provide effective feedback for real-time readjustment of key parameters (*44*). Overall, the interactive QCM capability encourages adaptive experimentation and reduces trial and error cycles, especially with biological samples to which access is limited.

As compared over currently available software solutions (*30, 32-34, 45-47*), QCM is parameter-free software package, requiring no parameters other than those characterizing the optical system; non-expert users can therefore easily operate it. Overall, QCM features an optimized software interface and display with easy-to-evaluate color-coded maps and histograms generated in real-time. This instantaneous quantitative information enables parameters to be adjusted, imaging conditions to be optimized and sub-regions of interest in the sample to be explored dynamically. Assessing such quality control of raw SMLM data at the earliest steps of acquisition enables an acceptance or rejection decision to be made on the basis of just a few hundred images, and thus optimizes the amount of data to be acquired, stored and analyzed for proper quantification of relevant observables.

It should be noted that the overall computation rate currently achievable here in real-time is mainly ensured by the UFUL module, which is based on a one-Gaussian fitting hypothesis, i.e., for low *D*_frame_ value, ideally below 1.0 particles/µm^2^/frame. It is therefore advisable to perform post-processing analyses of the recorded SMLM data, and to use dedicated algorithms to quantify any effective non-uniformly distributed molecules (*21, 23*). In this framework, we previously implemented UNLOC, a parameter-free algorithm approaching the Cramér-Rao bound for particles at high-density per frame and without any prior knowledge of their intensity (*35*).

We would like to underline that the QCM off-line mode offers invaluable possibilities to be used with post-acquisition SMLM data. We further stress that this mode is perfectly suited to carrying out standardized studies with no a priori assumptions on reusable SMLM raw metadata or during the review process of publications including SMLM data. QCM is also of general interest for teaching basic SMLM concepts to a wide audience. Overall, QCM can be seamlessly integrated into the workflows of homemade or commercial systems and cloud-based data analysis frameworks.

At present, we have succeeded in analyzing 2048 × 2048 pixels images at a rate of over 100 frames/second, a rate fast enough to explore dynamic processes in living samples. However, if it is possible to record data at a faster acquisition rate - for example, by focusing on a small ROI - we might face an intrinsic limitation of the SMLM technique due to the fact that the number of photons collected will be limiting at some point. Alternatives such as the promising event-based vision sensor-based imaging method (*48*) for *in vivo* imaging pave the way for a very promising paradigm shift in cell biology by giving access to a new quantitative set of relevant observables.

## MATERIALS and METHODS

### Ultra-Fast Unsupervised Localization (UFUL)

This section describes the mathematical basis of the Ultra-Fast Localization (UFUL) conception.

### Computational optimization of the GLRT detection

The detection step is based on previously mathematical developments (*35, 36*). In summary, when a particle is present, the PSF is modeled by a Gaussian *g*_*p*={*i,j*}_(*i*_0_, *j*_0_, *r*_*i*_, *r*_*j*_) centered in (*i*_0_, *j*_0_) ∈ ℝ^2^ and of dimension *r*_*i*_ and *r*_*j*_ :

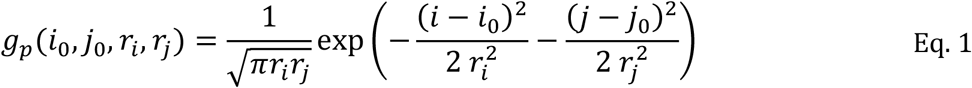

where the constant of normalization is such that ∬ *g*^2^ = 1.

The detection theory cannot estimate at the same time the value of the parameters (*i*_0_, *j*_0_, *r*_*i*_, *r*_*j*_) and the presence or absence of a particle (*49*). For the GLRT detector, the PSF is in the center of the window and *r*_*i*_ = *r*_*j*_ = *r* is known. Thus, the GLRT assesses the presence of a signal at each pixel such that (*i*_0_, *j*_0_) = (*i*_*n*_, *j*_*n*_)|_*n*∈ℕ_. When a signal is detected, the estimator searches in (*i*_*n*_, *j*_*n*_) the sub-pixel positions (*i*_0_, *j*_0_) of the PSF. For a PSF in a window, it is easier to write (*i*_*n*_, *j*_*n*_) = (0,0) for simplification purposes.

For a GLRT at known background (*35*), the mean *m* and variance *σ*^2^ of the background are known. This detector is based on the two *H*_0_ and *H*_1_ hypotheses, both perturbed by independent identically distributed additive Gaussian noise. For *H*_0_ in the working window *ω*, the signals at pixel *p* = {*i, j*} are the sum of background *m* and noise *n*_*p*_ of variance *σ*^2^:

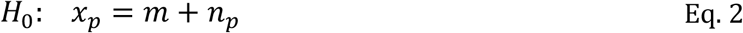

The *H*_1_ hypothesis has a Gaussian centered in the window that is modeled by:

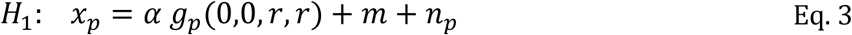

where *α* is the particle intensity.

Let *L*_0_ be log-likelihood of the *H*_0_ hypothesis:

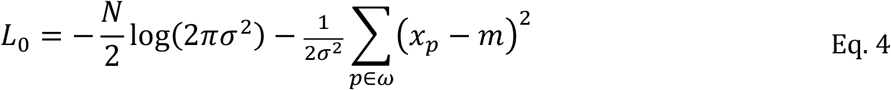

where *N* is the size of the window *ω*.

Let *L*_1_ be the generalized log-likelihood of *H*_1_ hypothesis:

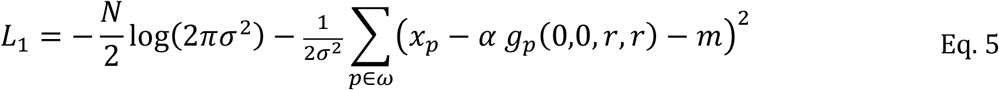

The estimated intensity is given by:

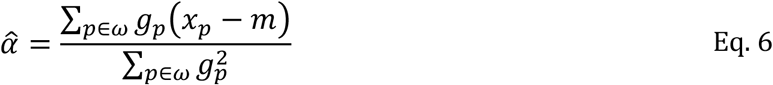

Thus, for a test based on the detection theory (*49*), the *H*_0_ hypothesis is rejected with a probability of false alarm *PFA* ∈] 0,1] if:

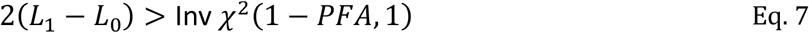

with Inv *χ*^2^(1 − *PFA*, 1) the inverse law of *χ* ^2^ with one degree of freedom. Thus, this test discriminates that, for a given *PFA*, the window contains either noise alone or a particle of *SNR* > 20 *dB*, with a detection probability *PD* ≈ 100% (*49*).

Here, we rewrite the GLRT expression to significantly optimize the computation time but without simplifying the robustness of the mathematical model.

Let *S*_*PFA*_ = Inv *χ*^2^(1 − *PFA*, 1) be the detection threshold, we can write the GLRT for a given pixel as:

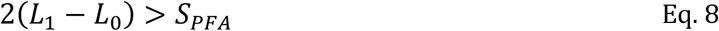

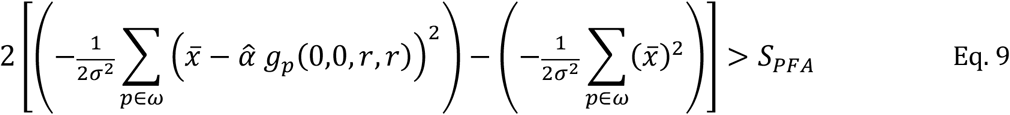

with 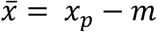.

Thus,

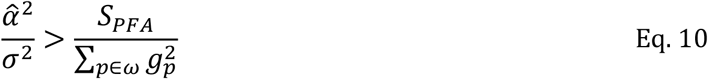

This requires first estimating the background mean 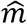 and variance 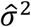 as previously described (see in (*35*) the note S6 in Supporting Material,). In practice, they are estimated once every 50 frames.

This test can therefore be performed for all pixels of a given frame. Computing the left term of Eq. 10 simply as a convolution (Eq. 6) provides the corresponding image of the *α* and GLRT values of the pixels. When the test is true in the region of interest (ROI), it corresponds to a particle defined as a single pixel or as a set of pixels for bright ones, from which a list of detected particles with an integer pixel value is established.

### Estimation of the particle localization

Once the particles are detected, the objective is to determine their subpixel localization, *i*.*e*., at which the signal intensity 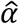 is maximum. Two computational methods are classically implemented:

- The ones based on an algorithm that performs oversampling of the 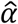 image are computationally expensive and cannot estimate the *r*_*i*_, *r*_*j*_ radii of the PSF;
- The others based on iterative fitting computation to estimate *r*_*i*_, *r*_*j*_, *i*_0_, *j*_0_ are also time consuming.

Here, we demonstrate that a third alternative is possible to determine the position of the particles and their radius with sub-pixel accuracy while guaranteeing an ultra-fast computational speed, meaning at a speed higher than that of image acquisition.

The PSF is modeled by a Gaussian and the algorithm is using the logarithm of *α* to obtain a quadratic expression. This enables a literal expression from which to derive the estimation of *r*_*i*_, *r*_*j*_, *i*_0_, *j*_0_, corresponding to the PSF sizes and sub-pixel coordinates of each particle, respectively.

As such, the current expression of 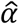 needs to be rewritten to provide a fast and efficient estimate of these parameters. By replacing *m* by its estimated value 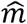, we obtain:

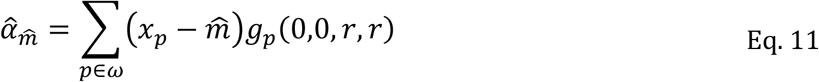

However, 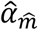 does not correspond to the minimal mean square error (MMSE) estimator that is expected from the solution given by 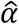. It is still possible to obtain a variance that coincides with that of the MMSE on the coordinates. When the *H*_1_ hypothesis is true (Eq. 3), we rewrite 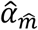 as:

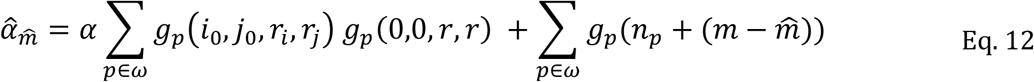

The second term 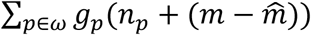 is a noise processed by a matched filter, but 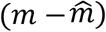, the noise term on the estimate of *m*, implied that 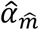 is a good approximation of a MMSE filter.

### Ultra-fast estimation of PSF dimensions and particle sub-pixel positions

We first detail the expression for estimating the intensity 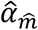 and the PSF sizes with logarithms. The characteristic PSF sizes are *r*_*i*_, *r*_*j*_ and *i*_0_, *j*_0_ are the sub-pixel positions of the particle. Moreover, PSF images modeled by Gaussians, filtered by a Gaussian kernel generate Gaussians. Thus, the intensity estimate is equal to:

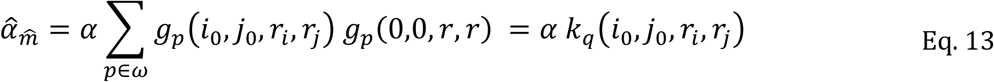

with 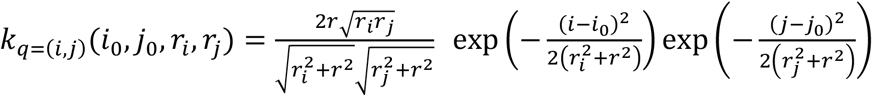.

The discrete second derivative on the *i*-axis of the logarithm of 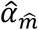 calculated at the positions (*i*_*n*_, *j*_*n*_) ∈ ℕ^2^ of the detected particles is:

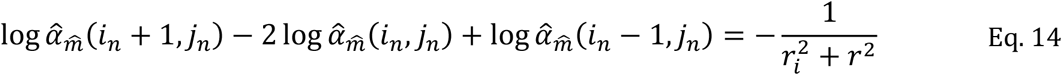

Thus, the estimator of the PSF sizes is:

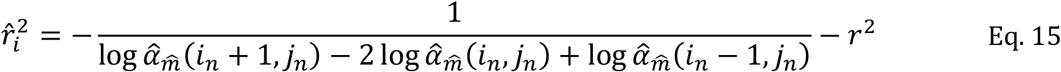

Furthermore, the discrete first derivative on the *i*-axis of the logarithm of 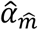 :

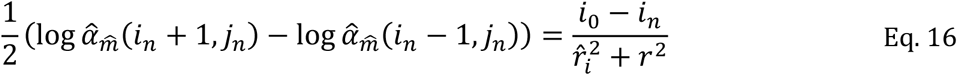

Then the estimator of *i*_0_ is:

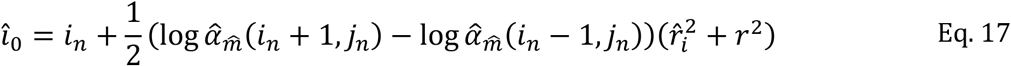

Similarly, for the *j*-axis, the estimators are:

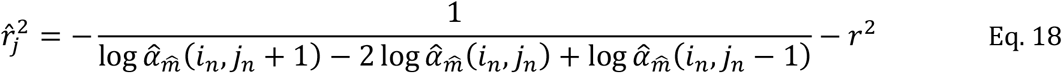

and

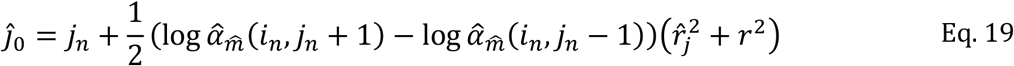

Thus, by computing only five logarithms of the image 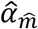, *i*.*e*., 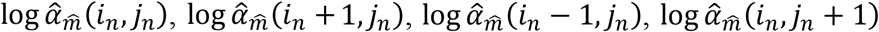 and 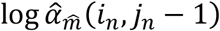, we can estimates all the parameters 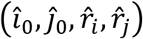 for the size and sub-pixel localization of a particle.

Then, in addition to calculate 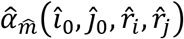 and the mean square error (MSE) to determine the SNR and variance of the error of the positions, it is necessary to calculate at the sub-pixel position, 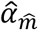 as well as the MSE on the error on these corresponding positions.

### Evaluation of the algorithm performances

All evaluations to compare the mathematical models or validates the algorithms were performed as shown on simulated images generated at a given SNR, PSF size, particle density/frame or image sizes. The codes used to generate these datasets are available on request from the authors.

For the evaluation of the CPU/GPU UFUL computation rate performances, the analyses were obtained with the following computer configuration: DELL Precision 7740 laptop; Central Processing Unit (CPU): E-2286M, 64 GB RAM; Graphics Processing Unit (GPU): NVIDIA Quadro RTX4000M. For CPU computations, the code is compiled in C for Matlab (MEX), using Advanced Vector Extensions (AVX) for 8-float 32-bit (single) operations to handle parallel computations. For GPU computations, the code is compiled with CUDA for Matlab (MEX-CUDA). The computation times correspond to the analysis of 16-bit RAW images stored in the PC RAM, from which the detection/estimation process provides the list of particles (position, size of 2D or 3D PSF astigmatism, measured intensity, SNR, noise level, position error) in the PC RAM.

Simulations used to demonstrate the real-time performances of QCM will be available under an approved open source license at the time of journal publication.

### Software and code

QCM is a multi-thread application developed on C/CUDA code on a LINUX platform (LINUX Ubuntu 20.04.2 LTS). It requires a CUDA toolkit for NVIDIA GPU. Two modes are available: a virtual mode for post-acquisition data evaluation that only requires a Graphic Programmable Unit (GPU) and an acquisition mode requiring in addition a PCO Edge 4.2 CLHS sCMOS camera. The data acquisition has been prioritized to offer the maximum frame rate of the camera (i.e., ≈ 100 frames per second). All other processes are running in parallel on specific and dedicated threads.

A QCM package is freely available online (see Supplementary Materials) for academic and nonprofit users. It includes a user guide, a set of experimental and synthetic data, and videos illustrating the visualization and quantification of observations.

### Data acquisition and analysis

All acquisitions were made using total internal reflection fluorescence (TIRF) illumination on a custom-built system based on an inverted microscope (Nikon, TE2000-U) as previously described (*35*) with a CFI Apo TIRF 100× NA 1.49 oil immersion objective (Nikon), a Argon/Krypton multiline laser (Innova 70C-Spectrum, Coherent Inc.), an axial drift correction by the autofocus module, except that the images are acquired with a PCO Edge 4.2 Camera link High Speed (CLHS) scientific Complementary Metal-Oxide-Semiconductor (sCMOS) camera on a LINUX platform. The microscope was controlled with a homemade Labview v.2021 (National Instruments) code, while the present homemade QCM code was used to acquire the data.

During acquisitions, data are evaluated in real-time with QCM to set appropriate acquisition conditions. Post-acquisition data analyses were performed with UNLOC (*35*) in high density mode with a high spatial frequency variation of background, a reconnection process with one Off-state lifetime frame and an integrated Gaussian rendering process after drift correction by correlation and without data filtering.

The raw experimental data illustrating the QCM performances are available on request from the authors.

### Reagents and sample preparations for experimental data

Quantitative experiments were performed with DNA-origami with GATTA-PAINT HiRes 80R nanorulers and the results evaluated using GATTAnalysis v1.5 software (GATTAquant).

COS-7 cells (ATCC CRL 1651™) and HeLa cells (ATCC CCL-2™) were grown in DMEM (Gibco) supplemented with 10% bovine fetal serum, 10 mM HEPES, 2 mM L-glutamine, 1 mM sodium pyruvate (Gibco), and 1% penicillin/streptomycin (Gibco).

For the dSTORM experiments, cells were plated on coverslips N° 1.5 of 18 mm diameter (Marienfeld GmbH, #0117580) and incubated at 37°C, 10% CO_2_ for 48 h before staining procedures. Cells were washed twice with pre-warmed (37°C) phosphate-buffered saline (PBS) before being fixed with 4% PFA into PBS for 15 min at room temperature (RT) than washed with PBS and treated with 50 mM NaBH_4_ for 10 min to reduce background fluorescence and finally washed with PBS. Fixed cells were permeabilized with 0.3% Triton™X-100 in PBS, for 30 min, then washed 3 times in PBS, and saturated with 3% bovine serum albumin (BSA) in PBS, for 45 min to reduce unspecific labeling. Nuclear pore complex protein Nup133 or β-tubulin were labeled overnight at 4°C with rabbit anti-Nup133 antibody (Abcam, #ab155990) and mouse anti-human β-tubulin mAb (Sigma-Aldrich, #05-661-I), respectively. Cells were then washed 5 times with 1% BSA in PBS before being incubated 30 min at room temperature with Alexa Fluor 647 AffiniPure™ Goat Anti-Rabbit IgG (H+L) (Jackson ImmunoResearch Europe Ltd., #111-605-003) for Nup133 labeling and Alexa Fluor™ 647-conjugated F(ab’)2-goat anti-mouse IgG (H+L) (ThermoFisher Scientific, #A-21237) for β-tubulin. After 5 washes with 1% BSA in PBS, cells were fixed again with 2% PFA, PBS for 5 min. Finally, after 3 washes in PBS, samples were mounted in depression slide with freshly prepared dSTORM buffer (50mM Tris, 50 mM NaCl a pH 8.0 supplemented with 50 mM cysteamine). Sealed samples with Twinsil® Speed 22 (Picodent, #1300 1002) were ready to be imaged.

For the two-color DNA-PAINT experiments, the HelaCells_Tubulin_Tom20 smart samples (Abbelight, France) were prepared in µ-slide VI 0.5 (IBIDI GmbH, #80607) using the microfluidic system Smart Flow (Abbelight, France) and the Smart Staining Kit instruction (Abbelight, France). Next, instructions for the DNA-PAINT kit (Massive Photonics GmbH, #MASSIVE-sdAB-FAST 1-PLEX) with anti-mouse and anti-rabbit nanobodies were applied to recognize the mouse anti-α-tubulin mAb (Sigma-Aldrich, #T6188) and rabbit recombinant anti-TOM20 mAb (abcam, #ab232589) antibodies, respectively. The α-tubulin and mitochondria were detected in a unique imaging buffer with a mix of imager 1 (Cy3b) at a final concentration of 0.5 nM, and imager 2 (ATTO 655) at the specified concentration, respectively. Sequential acquisition was performed with a stack of 50,000 frames recorded at 514 nm (200 mW) using a 525/50 filter for α-tubulin and a stack of 20,000 images was recorded at 647 nm (155 mW) using a 710/75 filter for TOM20.

## Acknowledgments

The authors thank Dr. Jérôme Touvier for his initial work on this project, Dr. Marc Allain (Institut Fresnel) for his deep reading of the UFUL part and Dr. Rémi Lasserre for general discussions.

## Author contributions

N.B., S.M., and D.M. conceived the project. N.B. and S.M. developed the algorithm, M.D. and R.F. performed the experiments to test extensively the software. D.M. supervised this work and prepared the original draft with the support of N.B. and S.M. All authors revised and edited the manuscript.

## Data and materials availability

The simulated and experimental datasets that illustrated the findings of this study are available from the corresponding authors upon request.

The QCM code is protected by the certificate Inter Deposit Digital Number (IDDN): IDDN.FR.001.510001.000.S.C.2022.000.31235 issued by the Agency for the Protection of Programs. The package is freely available online (see Supplementary Information) for academic and nonprofit users. It includes a user guide, a set of experimental and synthetic data, and videos illustrating the visualization and quantification of observations.

## SUPPLEMENTARY MATERIALS

### Supplementary Materials for this manuscript include the following

- A supplementary text for validation of the UFUL module and information on QCM software with figures S1 to S5.
- Movies S1 to S4

### Captions for movies S1 to S4

**Movie S1 – QCM user interface**

QCM requires only the setting three physical parameters (camera binning, exposure time and PSF size *r*_0_). QCM in real-time quality control indicators in the form of histograms (SMLM reconstruction, D_frame_, SNR, Precision, and estimated PSF size *r*_0_) and corresponding maps.

**Movie S2 -QCM performances on synthetic data**

QCM analysis is perfomed at 100 fps on 6,000 synthetic 2048 × 2048 pixel images with spatial densities ranging from 0.005 to 1.5 part/µm^2^/frame (see Suppl. Fig. 5). Histograms and maps of key indicators are updated instantly when the zoomed window is dragged to another areas of the image.

**Movie S3 - Adjustment of acquisition settings based on real-time QCM analyses**

Real-time QCM analysis of dSTORM acquisition parameters for β-tubulin imaging in COS-7 cells. Histograms and maps of key indicators are updated instantly when the zoomed window is dragged within the image. This allows a close inspection of different ROIs to adjust acquisition parameters in a few hundred images before starting acquisition.

**Movie S4 - QCM user interface for multi-color SMLM acquisition**

For multi-color SMLM acquistion, the QCM procedure is illustrated on a 256 × 256 pixel synthetic image dataset. Simulated objects of immunoglobulin-like shape are encoded in three particle types, which are sequentialy simulated with parameters specified as follow:

- chanel #1, 5 000 frames with PSF size *r*_0_ = 1.25 *pixels*, 20 ms exposure time, *D*_frame_ = 0.3 part/µm^2^/frame, SNR = 27 dB and a corresponding precision of 17 nm for red fluorescent particles;
- chanel #2, 10 000 frames with *r*_0_ = 1.15 *pixels*, 10 ms exposure time, *D*_frame_ = 0.1 part/µm^2^/frame, SNR = 30 dB and a corresponding precision of 13 nm for green fluorescent particles;
- chanel #3, 2 000 frames with *r*_0_ = 1.45 *pixels*, 15 ms exposure time, *D*_frame_ = 0.2 part/µm^2^/frame, SNR = 32 dB and a corresponding precision of 11 nm for bleu fluorescent particles.

