## Supplementary Materials for "QCM: real-time quantitative quality control of single-molecule localization microscopy acquisitions"

Sébastien MAILFERT *et al.*

**This PDF file includes:**

- Supplementary text for validation of the UFUL module and information on QCM software with figures S1 to S5.
- Legends for movies S1 to S4

**Other Supplementary Materials for this manuscript include the following:**

- Movies S1 to S4

*Distributed under a Creative Commons Attribution 4.0 International license.* A CC-BY-NC public copyright license has been applied by the authors to the present document and will be applied to all subsequent versions up to the Author Accepted Manuscript arising from this submission, in accordance with the grant's open access conditions.

### Supplementary Text

#### 1 Validation using synthetic data

##### *UFUL versus MMSE estimator performances as a function of the SNR*

The comparison was done on realistic synthetic data, *i.e.* on data close to the levels of noise, signal, PSF size, etc. that are typically the ones observed on experimental SMLM.

Thus, we introduce a signal-dependent noise:

$$x_p = \mathrm{N}\left(m_p, \sigma_p^2 = 2G\, m_p\right), \qquad \text{Eq. 1}$$

where $G$, the gain of an EMCCD[1] camera is equal to 1, and $m_p$, the average of $x_p$ is defined as

$$m_p = \alpha\, g_p(i_0, j_0, r_0, r_0) + m.$$

The PSF SNR ranges from 20 to 40 dB, a PSF size $r_0 = 1.2\ pixels$ and $m = 300$. We completed 1,000 noise draws, each containing a particle randomly localized in a square of 1-pixel side to compare the performance between UFUL and an MMSE estimator (Fig. S1).

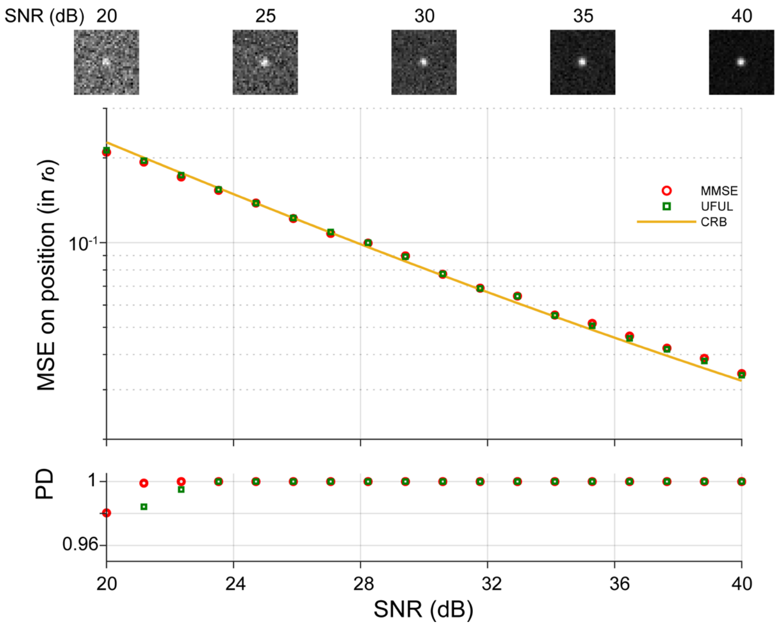


**Figure S1. Comparison between UFUL and an MMSE estimator as a function of SNR**

The MSE $\left(\sigma_{MSE} = \sqrt{\sigma_i^2 + \sigma_j^2}\right)$ on the determination of particle position are expressed in $r_0$ and the probability of detection $PD$ of the GLRT are plotted for different $SNR$. Thumbnails illustrate at a given $SNR$, one of 1000 simulated images performed with PSF size $r_0 = 1.2\ pixels$ and $m = 300$.

▶ The results obtained with UFUL overlap those of an MMSE estimator; both being close on the CRB. For a PFA of $10^{-6}$, the detection probability $PD \approx 100\%$ for any $SNR > 20\ dB$.

---

[1] The algorithm is robust and suitable for SMLM analyses acquired with EMCCD and sCMOS cameras, despite the fact that these two main types of cameras differ in electronics, pixel sizes, amplification modes or noises.

##### *Performance of the UFUL estimator as a function of PSF size and detection window*

Then, we compare the UFUL performances to an MMSE estimator as a function of PSF size on a synthetic dataset of 1,000 noise draws, with a SNR set to 30 dB and for UFUL, an adaptative filter size set to $r = 1.2\ pixels$ (Fig. S2)

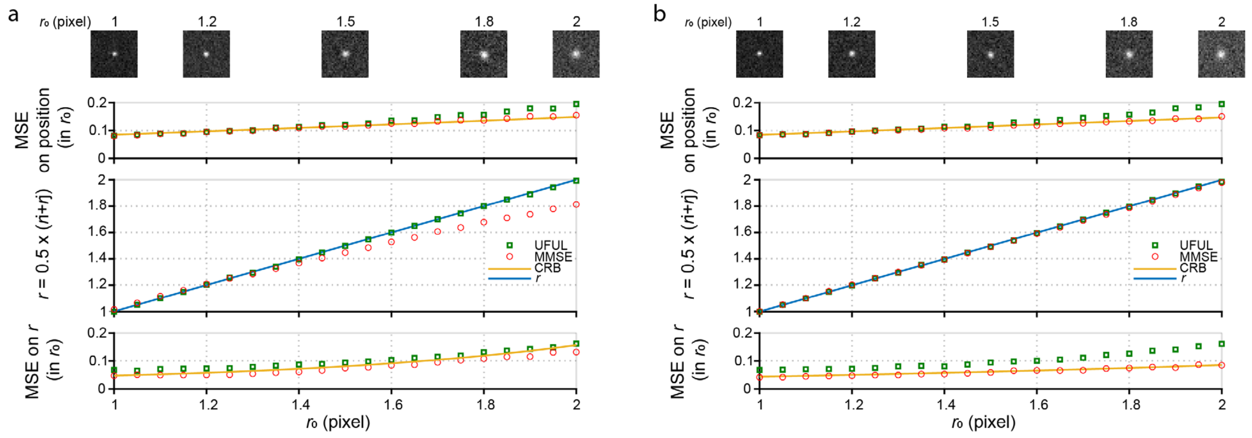


**Figure S2. Comparison between UFUL and an MMSE estimator as a function of PSF size**
The MSE $\left(\sigma_{MSE} = \sqrt{\sigma_i^2 + \sigma_j^2}\right)$ on the determination of particle position expressed in $r_0$ and the probability of GLRT detection are plotted for different SNR. The thumbnails illustrate a random draw for different SNR. Simulations were performed with PSF size $r_0 = 1.2\ pixels$ and $m = 300$. a) $\omega = 8 \times 8$, b) $\omega = 15 \times 15$.

For a working window of $\omega = 8 \times 8$ (Fig. S4a), the PSF size, ranging from $r_0 = 1.0\ to\ 2.0$, allows estimation of $i_0, j_0, r_0$ but with a slight bias on $r_0 > 1.6$, still with a variance $\sigma_{\hat{r}_0}$ slightly above the CRB; the small size of the working window biases the MMSE results. For UFUL, the variance in particle positions is comparable to that of MMSE and it is close to the CRB considering that the adapted filter of UFUL is only accurate on the synthetic data where $r = r_0 = 1.2$. Similar analyses were performed for a working window of $\omega = 15 \times 15$ (Fig. S4b). In that case, the results of MMSE estimator have little bias and the variance of $\hat{r}_0$ is very close to the CRB.

▶ In conclusion, UFUL provides, with an accuracy close to the CRB, the estimation of $r_0$ on both $i$ and $j$ axes with the estimator $\hat{r}_0 = (\hat{r}_i + \hat{r}_j)/2$ regardless of the PSF size and the working window.

##### *UFUL performances to evaluate PSF under 3D astigmatism conditions*

Taking advantage of the fact that UFUL enables the estimation of the PSF on both $i$ and $j$ axes, we, therefore, tested its performance to determine the particle localization as a function of their optical axial position around $z_0$, the focal point. This was done on synthetic data mimicking image acquisition with an astigmatic lens (*1*). Here, the PSF size $(r_i, r_j)$ varies on $i$ and $j$ axes in pixels from (1.2, 3.0) to (3.0, 1.2). Again, we generate a synthetic dataset consisting of 1,000 noise draws, with an SNR set to 30 dB and for UFUL, an adaptative filter size set to $r = 1.2\, pixels$. As illustrated below (Fig. S3), the PSF sizes are well estimated with a slight bias around 3.0 pixels.

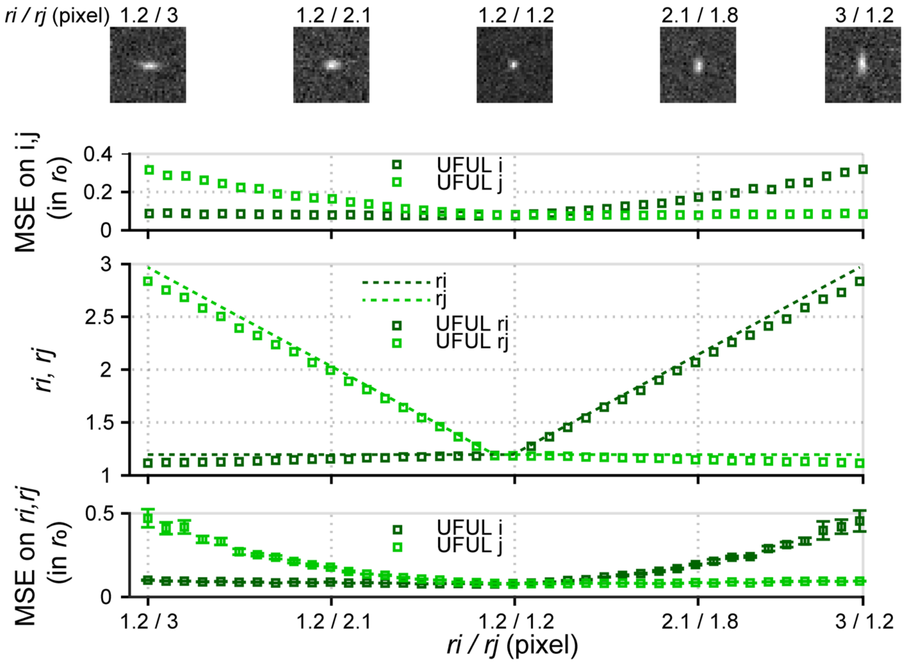


**Figure S3. UFUL performance for estimating PSF positions and sizes in 3D astigmatism conditions**

The RMS error on the PSF position (upper plot), the PSF size estimate (middle plot) and its RMS error (lower plot) are determined for the $i$ and $j$ axes, respectively, as a function of the PSF deformation. The thumbnails illustrate random draws for given $r_i/r_j$ratios.

#### 2 Quality Control Map algorithm

As previously stated, the UFUL performances allow designing an algorithm that processes the UFUL output (particle positions, PSF size, intensity, and background) to provide quality control indicators (signal-to-noise ratio, density, precision maps, and histograms).

##### *Calibration of the density $D_{frame}$*

As no algorithm can detect all particles, especially those that cannot be resolved because they appear in too close proximity to each other in a single frame, the density can be poorly estimated. Thus, a calibration of $D_{\text{frame}}$ is necessary, as UFUL can be biased in its estimation, since it was designed for low density data, i.e., below 2.0 particles/µm²/frame at usual SNRs (*2*). Indeed, above this $D_{\text{frame}}$ threshold, the percentage of non-resolved particles becomes critical and significantly alters the accuracy of particle localization, even for those with a high SNR. A calibration was been therefore carried out on synthetic data, at different densities and for a known $r_0$ (i.e., $r_0 =$

$1.25\,pixels$) (Fig. S4). In practice, this calibration has been integrated so that QCM returns indicators with realistic density values.

The values of the "uncalibrated D" analysis, $D_{\text{uncal}}$ are obtained by QCM without prior correction. This curve is approximated by a polynomial of order 2: $f_{1.25}(x)$ and the calibration obtained by $D_{cal} = f_{1,25}^{-1}(D_{uncal})$. Thus, it is possible to obtain a curve invariant to $r_0$ as follow:

$$D_{cal} = \left(\frac{r_0}{1,25}\right)^2 f_{1,25}^{-1}(D_{uncal}) \qquad \text{Eq. 2}$$

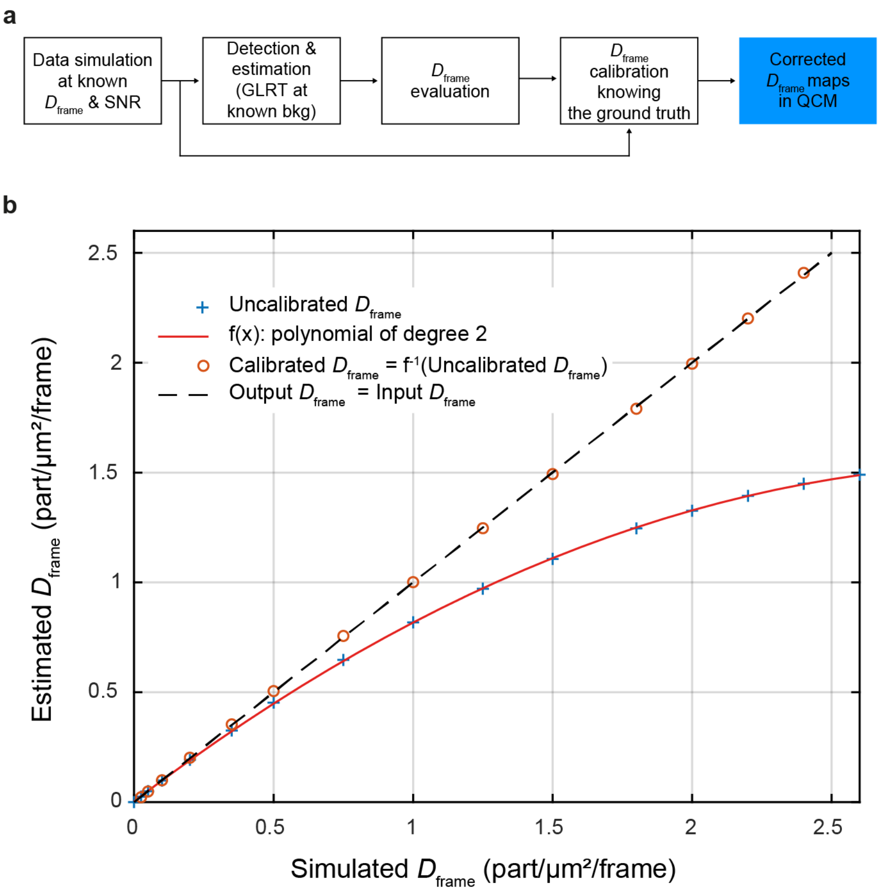


**Figure S4. Calibration of the density evaluation**

a) Workflow of density calibration. Data were simulated at known density and signal-to-noise ratio. The estimated density is compared to the simulation and corrected to correspond to the simulated one. QCM displays the corrected densities. b) Density correction curves. The x-axis corresponds to the density value of the synthetic data; the y-axis to the output of QCM expressed in part/µm²/frame. Evaluated density (blue cross) results from an underestimation of the real density. A fit with a quadratic function (red curve) matches the estimated density after correction with the simulated density.

▶ In practice, this calibration is integrated so that the algorithm returns indicators with realistic density values.

#### *Practical features*

Based on a simple graphical user interface (Fig. 4a), QCM is an effective unsupervised algorithm that requires setting the following parameters:

- $r_0$, the characteristic PSF size of the optical setup according to the full width at half maximum. This value must be expressed in pixels. Note that QCM displays the current $r_0$ PSF size evaluated during the data acquisition.
- the binning and exposure time parameters of the camera.

Histograms of estimated $r_0$, $D_{\text{frame}}$, SNR, and the precision of particle localization are displayed in real-time with the option to display an SNR-$D_{\text{frame}}$ space diagram representation (*2*) for the full frame, zoomed region and for the entire stack or only the last frames. The corresponding maps are also computed and displayed in real-time.

QCM offer the additional advantage of estimating key quality control indicators for the full frame or the zoomed region (Fig. S5).

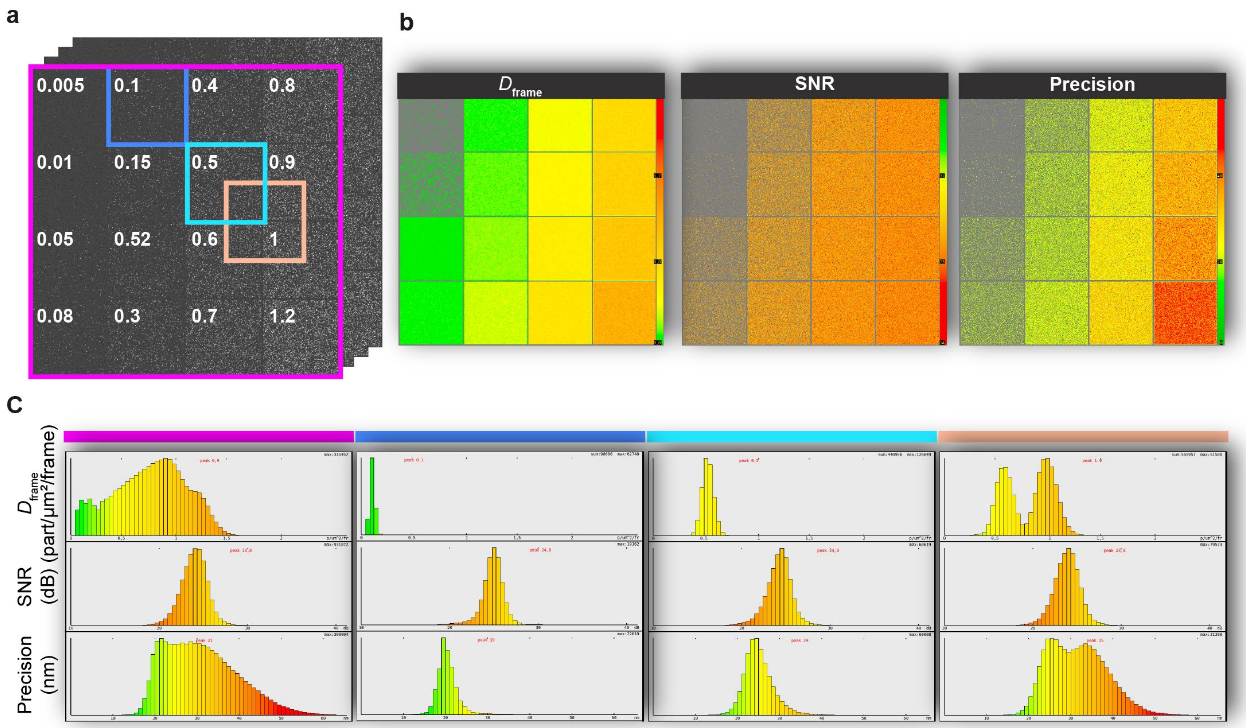


**Figure S5 – Local and global key quality control indicators**

a) 500 images of raw super-resolution data with constant PSF size ($r_0 = 1.25\ pixels$), constant SNR (25 dB) and different $D_{\text{frame}}$ ranging from 0.005 to 1.2 particles/µm²/frame. b) QCM map estimation of density, SNR and precision, respectively. c) The different ROIs shown in a) are updated in real-time allowing the quality control evaluation to be refined, especially for samples with an inhomogeneous distribution of the molecule of interest.

***Software package***

The QCM software package contains:

- The QCM algorithm;
- A user manual with installation instructions;
- Simulated and real SMLM raw data images.

QCM will be freely available for academic and nonprofit use as a software package here which includes updated versions at the time of journal publication. The source code is available upon a signed material transfer agreement.

**CAPTIONS for MOVIES S1 to S4**

**Movie S1 – QCM user interface**

QCM requires only the setting three physical parameters (camera binning, exposure time and PSF size $r_0$). QCM in real-time quality control indicators in the form of histograms (SMLM reconstruction, $D_{frame}$, SNR, Precision, and estimated PSF size $\hat{r}_0$) and corresponding maps.

**Movie S2 - QCM performances on synthetic data**

QCM analysis is perfomed at 100 fps on 6,000 synthetic 2048 × 2048 pixel images with spatial densities ranging from 0.005 to 1.5 part/µm²/frame (see Suppl. Fig. 5). Histograms and maps of key indicators are updated instantly when the zoomed window is dragged to another areas of the image.

**Movie S3 - Adjustment of acquisition settings based on real-time QCM analyses**

Real-time QCM analysis of dSTORM acquisition parameters for β-tubulin imaging in COS-7 cells. Histograms and maps of key indicators are updated instantly when the zoomed window is dragged within the image. This allows a close inspection of different ROIs to adjust acquisition parameters in a few hundred images before starting acquisition.

**Movie S4 - QCM user interface for multi-color SMLM acquisition**

For multi-color SMLM acquistion, the QCM procedure is illustrated on a synthetic 256 × 256 pixel image dataset in which three particle types are sequentialy simulated with parameters specified as follow:

- chanel #1, 5 000 frames with PSF size $r_0 = 1.25\, pixels$, 20 ms exposure time, $D_{\text{frame}} = 0.3\ \text{part/µm}^2\text{/frame}$, $\text{SNR} = 27\ \text{dB}$ and a corresponding precision of 17 nm for red fluorescent particles;
- chanel #2, 10 000 frames with $r_0 = 1.15\, pixels$, 10 ms exposure time, $D_{\text{frame}} = 0.1\ \text{part/µm}^2\text{/frame}$, $\text{SNR} = 30\ \text{dB}$ and a corresponding precision of 13 nm for green fluorescent particles;
- chanel #3, 2 000 frames with $r_0 = 1.45\, pixels$, 15 ms exposure time, $D_{\text{frame}} = 0.2\ \text{part/µm}^2\text{/frame}$, $\text{SNR} = 32\ \text{dB}$ and a corresponding precision of 11 nm for bleu fluorescent particles.
